# A comparison of microglial morphological complexity in adult mouse brain samples using 2-dimensional and 3-dimensional image analysis tools

**DOI:** 10.1101/2024.09.30.615842

**Authors:** Colin J. Murray, Eva D. Tunderman, Haley A. Vecchiarelli, Fernando González Ibáñez, Marie-Ève Tremblay

## Abstract

Characterizing cell morphology has been an essential aspect of neuroscience for over a century to provide essential insights into cellular function and dysfunction. Microglia, the resident innate immune cells of the central nervous system, undergo drastic changes in morphology in response to various stimuli, with many classifications proposed in recent years. Increased availability of advanced analysis software to study microglial morphology represents a step forward in the field. However, whether the use of advanced analysis tools provides equivalent or varied outcomes remains undetermined. This study re-analyzed raw data, previously processed using a standard 2D microglial morphology analysis method, using 3D analysis methods. The published article observed significant changes in microglial morphology in the mouse ventral hippocampus after administration of a ketogenic diet and exposure to repeated social defeat stress in young adult male mice. Overall, we observed different statistical outcomes in the 3D dataset compared to the previously published 2D results, with both maintained and new findings. This may indicate that the 3D analysis method is better able to capture minute changes in morphology. However, overall conclusions on microglial morphology changes remain consistent between methods. Lastly, we highlight the difference between a nested statistical design, which considers within animal variability, and a non-nested design. Overall, we highlight and discuss differences between 2D and 3D microglial morphology analysis and explore the contribution of individual cell and animal variability to statistical outcomes.

**Main Points:**

- 3D analysis method generates similar and novel results compared to a 2D method.
- verall microglial morphological changes to stimuli are comparable between methods.
- nested statistical design produces distinct significant differences.

## Introduction

Microglia, the resident innate immune cells of the central nervous system (CNS), are critical to maintain CNS homeostasis and perform immune surveillance (Kreutzberg, 1996; Šimončičová et al., 2022). In healthy conditions, microglia constantly communicate with neurons and other cells of the CNS, promoting neuronal and glial cell health by clearing debris, pruning synapses, modulating inflammation, facilitating neuronal connectivity and plasticity, and neuronal/glial survival (Colonna & Butovsky, 2017; Gao et al., 2023). These essential functions are generally accompanied by changes in microglial morphology, with healthy adult grey matter microglia often exhibiting small somas with highly motile, ramified processes (i.e., surveillant morphotype) that enable them to interact dynamically with their microenvironment, as observed in adult mouse neocortex (Nimmerjahn et al., 2005; Tremblay et al., 2010). However, microglia in white matter or early life stages, for example, can display different morphologies in rodent models and humans, reflecting their distinct environmentally-induced functional states (Ambrose et al., 2020; Paolicelli et al., 2022; Tremblay, 2020; Vidal-Itriago et al., 2022). Upon detecting pathogens or signs of damage and debris, microglia can shift toward a reactive pro-inflammatory state, triggered by pathogen-associated molecular patterns (PAMPs) and damage-associated molecular patterns (DAMPs) via pattern recognition receptors (PRRs), like Toll-like receptors (TLRs) (Colonna & Butovsky, 2017). In response to such adverse signals, microglia undergo a variety of morphological changes, often retracting and thickening their processes, presenting somatic swelling, and sometimes becoming hyper-ramified. Indeed, the existence of mixed morphological states that are distinct from the surveillant morphotype of homeostatic microglia in health and disease and the diversity within these populations is often overlooked. In line with this, microglial reactivity and morphological changes are not binary, but exist as a spectrum of morphologies, each signifying different microglial states that often reflect and shape the local environment they inhabit (Fig. 1). Understanding this diversity is important. Even though morphology is not a direct measure of function, there is substantial evidence linking specific microglial morphological states with distinct transcriptomic profiles, for instance, suggesting that these morphological changes have significant functional implications (Gyoneva et al., 2019; Paolicelli et al., 2022).

**Figure 1.**
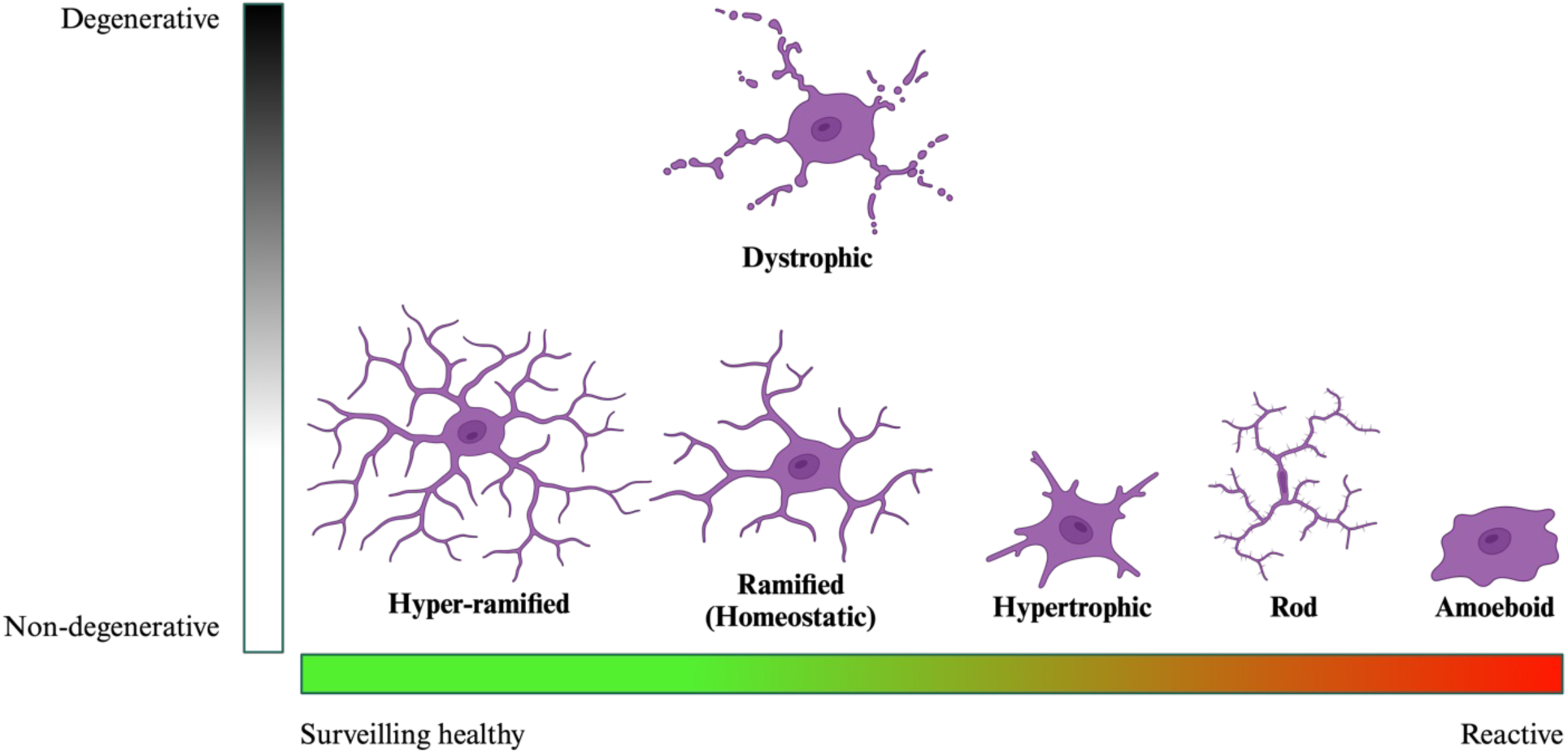
|. Microglia exist within a continuum in which their transcriptional and morphological characteristics often mirror the context of their surrounding microenvironment. These states, including ramified, bushy, and amoeboid morphologies, are dynamic and can change throughout the lifespan or in response to various stimuli. While individual cells may show variability within a single state, an overall shift in microglial morphology can occur depending on specific environmental or pathological signals. Created with Biorender.com.

For example, upon detecting PAMPs and/or DAMPs caused by infection, injury, apoptotic cells and/or cellular debris, microglia can retract their branches, undergo somatic swelling, and adopt a less complex structure, which is often associated with increased phagocytic activity (Colonna & Butovsky, 2017; B. M. Davis et al., 2017, 2017; Doorn, Moors, et al., 2014; Ginhoux et al., 2010; Morrison et al., 2017; Savage et al., 2019). During traumatic brain injury, neurodegenerative disease, or chronic inflammation, microglia can further retract their branches and adopt an amoeboid morphology, which is associated with enhanced migration efficiency to sites of damage and phagocytic activity in the affected areas (Doorn, Goudriaan, et al., 2014; Wicks et al., 2022). This ameboid microglia morphology is frequently observed in cases of severe or chronic inflammation, but also during healthy states, such as during development when microglial migration and phagocytosis are prevalent (Badanjak et al., 2021; Doorn, Moors, et al., 2014; Perez-Pouchoulen et al., 2015; VanRyzin et al., 2019). Another distinct morphology that inhabits the far-end of the spectrum is the hyper-ramified microglia, which has been observed to be more prominent in stress-resilient mice, indicating a potential link between this morphology and an adaptive response to stress (Fujikawa & Jinno, 2022). In addition, there is diversity within these contexts as well, with many intermediate or more polarized forms also being present, such as bushy microglia—characterized by a large soma and surrounded by short, stubby branches (Wicks et al., 2022). For a more extensive list of microglia morphologies, please refer to Table 1.

**Table 1.** |. Prominent examples of microglial morphologies and their associated significance.

| **Morphology** | **Characteristics** | **Functions** | **Associated Conditions** | **Model** | **References** |
| --- | --- | --- | --- | --- | --- |
| **Hyper-ramified** | Highly ramified, with complex and branched branches; appear elongated and thin | Adaptive response to acute and chronic stress; surveillance; potentially neuroprotective | Observed in stress-resilient mice | Mouse; rat | (Fujikawa & Jinno, 2022; Hellwig et al., 2016; Hinwood et al., 2012, 2013; Smith et al., 2019) |
| **Ramified**<br>**(Homeostatic)** | Small soma with long, highly branched branches actively surveilling the brain environment | Surveillance of CNS environment by extending and retracting thin ramified branches; detection of damage or pathogen associated patterns; maintenance of homeostasis | Healthy brain; baseline condition | Human; mouse; rat; zebrafish | (Colonna & Butovsky, 2017; Davalos et al., 2005, 2005; Ginhoux et al., 2010; Nimmerjahn et al., 2005; Svahn et al., 2013; Torres-Platas et al., 2014; Tremblay et al., 2010; Wake et al., 2009) |
| **Hypertrophic**<br>**(Bushy)** | Enlarged soma with increased ramification; shorter and thicker branches with increased complexity | Early response to injury; may facilitate the transition to more immune activated states | Initial injury or inflammation response | Human; mouse; rat | (Ambrose et al., 2020; Jensen et al., 1994; Ladeby et al., 2005; Wicks et al., 2022) |
| **Rod** | Straight, rod-shaped appearance; aligned branches; often observed near damaged neurons | Involved in response to axonal damage; potential guidance and repair functions; may interact directly with neurons | Acute injury; CNS infections (e.g. encephalitis); aging; neurodegeneration | Human; mouse; rat | (Bachstetter et al., 2017; Giordano et al., 2021; Holloway et al., 2019; Taylor et al., 2014; Yuan et al., 2015; Ziebell et al., 2012) |
| **Amoeboid** | Enlarged soma with very few or no branches; appears rounded and amoeboid-like | Highly phagocytic; involved in clearing debris and dead cells | Developing brain; clearing debris and apoptotic cells; acute injury response; demyelination conditions | Human; mouse; rat; zebrafish | (E. J. Davis et al., 1994; Kettenmann et al., 2011, 2013; Kreutzberg, 1996; Lawson et al., 1990; Morsch et al., 2015; Schwarz et al., 2012; Stence et al., 2001; Svahn et al., 2013) |
| **Dystrophic** | Fragmented swollen branches; reduced branching and phagocytic | Impaired phagocytic function, may contribute to | Neurodegenerative disorders (e.g., Alzheimer's, Parkinson's); |  | (Ritzel et al., 2019; Shahidehpour et al., 2021) |
|  | activity; senescence<br>marker-positive | neurodegeneration<br>and impaired tissue<br>repair; potentially a<br>form of microglia<br>senescence | aging; traumatic<br>brain injury |  |  |

In order to quantify these distinct microglial morphotypes, a suite of parameters has been developing since Pío del Río Hortega first discovered microglia in 1919. Some of the most commonly used parameters are soma area/volume, convex hull, circularity, branch length, Sholl analysis, and more (Leyh et al., 2021). Although each descriptor can give useful information as to how microglia are responding to any given stimuli, combinations of parameters in relation to one another can give increased insight into individual morphotypes. For example, the morphology index provides a readout of overall cell shape based on the ratio between soma area and arborization area (i.e., the larger the value, the greater the soma size in relation to branching area), whereas the Schoenen ramification index provides a readout of the overall branching complexity of the cell, represented by the ratio between the maximum number of Sholl intersections and the number of primary branches of the cell (Schoenen, 1982; Tremblay et al., 2012).

Given the association between microglial morphology and environmental pressures and/or stimuli, microglial morphology analysis is a crucial tool for investigating their functional state in CNS pathology. Techniques commonly used to assess microglial morphology often involve either two-dimensional (2D) maximum intensity projection analysis (Karperien et al., 2013; Kozlowski & Weimer, 2012; Torres-Platas et al., 2014; Verdonk et al., 2016) or more advanced three-dimensional (3D) reconstruction analysis methods (using software such as Neurolucida or Imaris) (Erny et al., 2015; Perego et al., 2013). However, it remains unclear whether these differing approaches yield comparable results and whether the conclusions drawn from each method are equally reliable and accurate. The potential discrepancies between these methods highlight a broader, increasing issue within the field of neuroscience: the lack of standardization in data generation, collection, and analysis, which significantly contributes to reproducibility challenges and translational failures (Green et al., 2022).

To this end, using Imaris v10.0.2, this study re-analyzed raw data in 3D from a previously published study from our group that revealed significant changes in 2D microglia morphology in the mouse ventral hippocampus after administration of a ketogenic diet (KD) or control diet (CD) and exposure to repeated social defeat stress in young adult male mice. The original analysis focused on comparing key morphological parameters, including microglial soma area, arborization area, and the morphology index, across different experimental groups. These groups comprised a control group not exposed to social defeat stress, stress-susceptible (SUS) and stress-resistant (RES) mice, further categorized based on their diet (KD *vs* CD). The 2D analysis highlighted distinct changes in microglial morphology, such as differences in soma size and branching complexity. By extending this analysis into 3D, using the same dataset, we aim to provide a comprehensive understanding of the differences between 2D and 3D morphology analysis methods.

## Methods

The raw data utilized in this methods manuscript were obtained from those published previously (González Ibáñez et al., 2023). Therefore, we only outline aspects of the method section necessary to contextualize these results.

### Animals

All animal experiments were approved by the Université Laval’s animal ethics committee (Comité de protection des animaux de l’Université Laval), in accordance with the guidelines set forth by the Canadian Council on Animal Care. For in depth details concerning animal experiments and the protocols used for the ketogenic diet, social defeat paradigm, social interaction test to determine resistant *vs* susceptible mice to stress, euthanasia, and tissue preparation, please refer to González Ibáñez et al. (2023). Briefly, 7–8-week-old male C57BL/6J mice (Jackson Laboratories) were perfused with phosphate-buffered saline (PBS; 50 mM pH 7.4) followed by a mixture of 4% paraformaldehyde and 0.2% glutaraldehyde in phosphate buffer (PB; 100 mM, pH 7.4) (González Ibáñez et al., 2023).

### Fluorescence Staining and Confocal Imaging

For a detailed description of the methods used for immunofluorescence staining and confocal imaging, please refer to González Ibáñez et al. (2023). Briefly, 50 μm coronal sections were incubated with sodium citrate at 70 °C, 0.1% NaBH_4,_ and blocking buffer before overnight incubation at 4 °C with mouse anti-ionized calcium-binding adapter molecule 1 (IBA1; EMD-Millipore cat# MABN92+; 1:150) and rabbit anti-transmembrane protein 119 (TMEM119; Abcam cat# ab209064; 1:300) primary antibodies. The following day, the sections were incubated with corresponding fluorescent secondary antibodies for 90 min and then with DAPI (1:20k) for 5 min. The sections were then mounted and coverslipped. Microglia (IBA1^+^/TMEM119^+^) (n=3–5 mice/group; 32–94 microglia/group) were then imaged at 40x with a Quorum WaveFX spinning disc confocal microscope (Quorum Technologies, Guelph, ON, Canada) with an ORCA-R2 camera (512 x 512 pixels; Hamamatsu Photonics, Hamamatsu, Japan). Z-stack images were taken with a Z-interval of 0.33 μm.

### Microglial Morphology

This study utilized confocal images from a previously published paper from our research group— González Ibáñez et al. (2023)—to reanalyze previous 2D data with the 3D morphological analysis software, Imaris (v10.0.2 Oxford Instruments). All microglial analysis was conducted blind to experimental conditions.

The following protocol was utilized to reconstruct microglia within the Imaris workspace for morphological characterization:

1. Raw czi. image files were imported into the Imaris ‘Arena’ and converted to the ims. file format using the built in Imaris file converter feature.
2. Once the image was selected, a new surface was created and the ‘Skip Automatic Creation, Edit Manually’ button was selected for manual reconstruction of the soma.

a. Under the ‘Contour’ tab, the ‘Time’ drawing mode with an Insert Vertex every 300 ms was used to trace the soma (DAPI^+^/IBA1^+^). The perimeter of the soma was traced every 2–5 slices throughout the Z distance of the soma, which proved to accurately represent the volume of the soma qualitatively.
b. The ‘Create Surface’ button was then selected, which allows Imaris to interpolate the slices not traced based off of the slices that were traced, completing the soma reconstruction. (Fig. 2B)
3. Next, we added another object using the ‘Add New Filaments’ selection for reconstruction of the microglial branches.
4. We then selected the ‘Skip Automatic Creation, Edit Manually’ button, opened the ‘Edit’ tab and selected ‘Import Soma…’ under the ‘Process Filaments’ section.

a. Our manual soma reconstruction object was selected.
5. We then opened the ‘Creation’ tab, checked the ‘Keep Data’ box and proceeded to recompute the filaments.

a. The ‘Tree Autopath Algorithm’ was used with default creation parameters.
b. The source channel was selected and the ‘Use Existing Segment’ option was selected under the ‘Starting Points’ heading.
c. Soma model was calculated and subsequently used as the starting point for filament tracing.
d. The thinnest filament diameter was set at 1 μm.
e. The seed point threshold was manually adjusted so that all visible branches extending from the soma contained at least one seed point at the base and terminal end. The ‘Classify Seed Points’ option was selected.
f. For seed points classification, a portion of seed points were manually selected or discarded before selecting ‘Train and Predict’—an option within Imaris that uses machine learning to select or discard seed points based off of your previous selections.
g. No ‘Terminal Segment Postfilter’ was selected.
h. Filaments were then inspected and manually edited if needed. (Fig. 2C)
6. We then utilized the ‘Filament Sholl Analysis24’ plugin, acquired from Ironhorse1618 (2023): https://github.com/Ironhorse1618/Python3.7-Imaris-XTensions.

a. Sholl Sphere Radius was set to 5 μm.
b. ‘Create Convex Hull surface’ was selected. (Fig. 2D)
7. Finally, under the statistics tab, the volume of the convex hull, volume of the soma, total Sholl intersections and critical radius were extracted.

**Figure 2.**
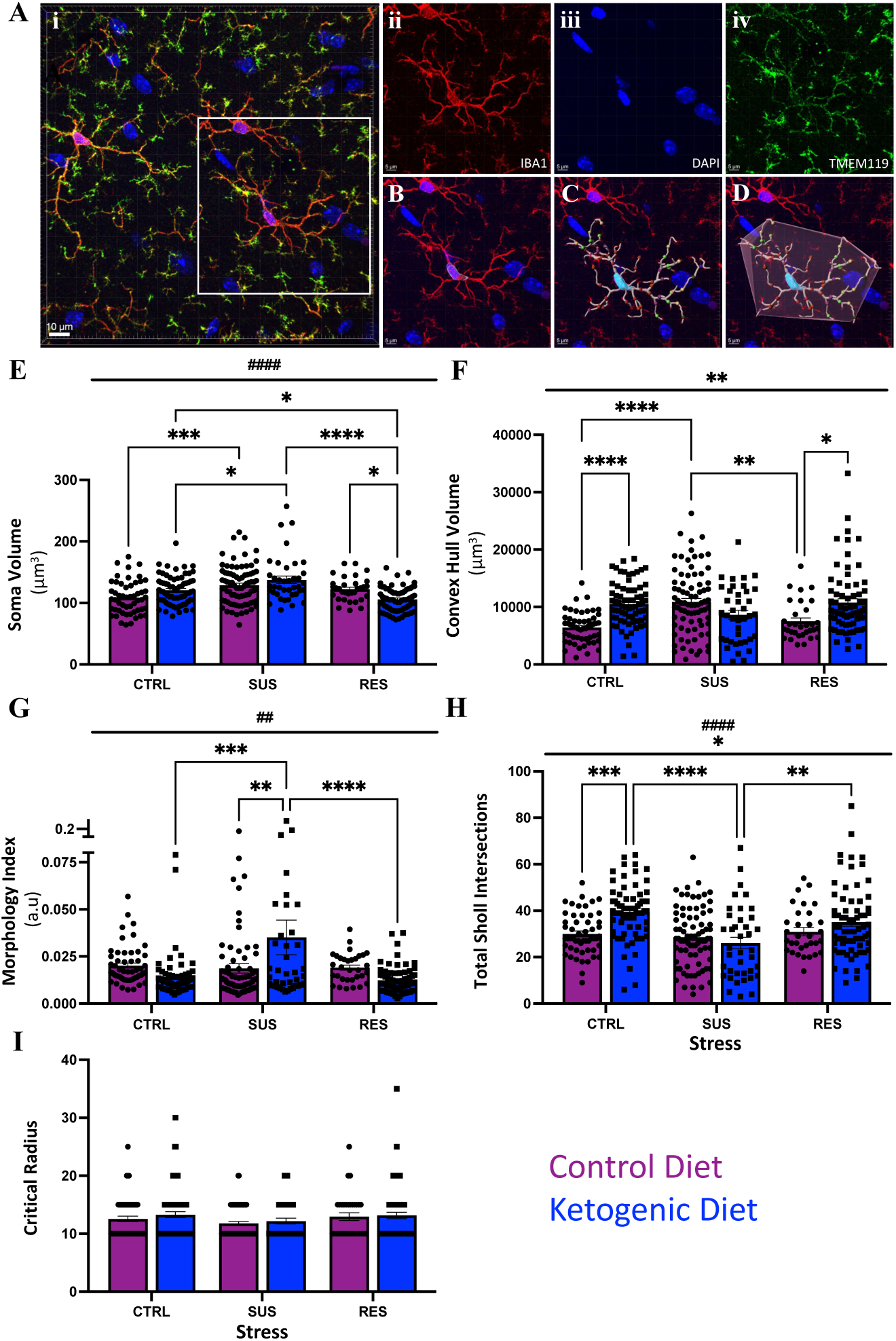
| Representative images of microglia (Ai) stained with immunofluorescent antibodies against IBA1 (red: Aii), TMEM119 (green: Aiv) and DAPI (blue: Aiii) visualized in the Imaris workspace. Soma reconstruction (B) in Imaris is shown alongside filament tracing and sholl analysis (C) and convex hull (D). Microglial morphological parameters from 3D analysis methods are shown (E–I). Microglia show significant adaptations in all morphological parameters, except for critical radius, in response to diet and stress (E–I). n=3–5 mice/group; n=32–94 microglia/group. Statistical significance was assessed using a 2-way ANOVA with Tukey *post hoc* test, and data are expressed as mean ± standard error of the mean, where \**p*≤0.05, \*\**p*<0.01, \*\*\**p*<0.001, *\*\*\*\*p*<0.0001. In addition, #≤0.05, ##<0.01, ###<0.001 depicts a main effect of stress. CTRL: non-stressed controls; SUS: susceptible to stress; RES: resistant to stress; a.u: arbitrary units.

Although many pre-processing options exist in Imaris, we chose not to use these in order to keep the images as similar as possible to those used for the 2D morphology analysis with the aim to identify if the 3D aspect of this analysis alters results.

For morphology, we analyzed soma volume (μm^3^), convex hull volume (μm^3^), and morphology index (soma volume/convex hull volume) for the 3D analysis method, which respectively compares with soma area (μm^2^), arborization area (μm^2^), and morphology index (soma area/arborization area) quantified using the 2D analysis method performed by González Ibáñez et al. (2023). In addition, we quantified the total number of Sholl intersections at 5 μm radius intervals and the critical radius (radius with the most Sholl intersections) for microglia using the 3D analysis method.

### Statistics

Statistical testing was performed in an identical manner as in González Ibáñez et al. (2023) to allow for comparison. However, due to the nature of the 3D analysis, some images that contained smaller Z-stacks and therefore had cut-off branches in the Z-plane previously used for the 2D analysis, could not be used. In addition, some images had multiple microglia analyzed where 2 or more fully intact microglia were present. Therefore, the sample size for the current study is slightly different (n=3–5 mice/group; 32–94 microglia/group) compared to González Ibáñez et al. (2023) (n=3–5 mice/group; 51–88 microglia/group).

Data shown are reported as means ± standard error of the mean (SEM). Normality of the data was assessed using the Shapiro-Wilk test. The data did not pass the normality test, but due to the fact that no non-parametric test yet exists for our study analysis design, parametric tests were used.

We first sought to perform statistical tests identical to those performed by González Ibáñez et al. (2023) to be able to compare our 3D analysis with their 2D data. This was accomplished by performing a 2-way analysis of variance (ANOVA) test with diet and stress as our independent factors. Our *a priori* hypothesis was that we would find similar significant results to González Ibáñez et al. (2023). To test this hypothesis, we also performed Tukey’s *post-hoc* test to assess these comparisons, as did González Ibáñez et al. (2023). This analysis was performed using GraphPad Prism (v10.0.3 Graphstats Technologies).

To directly compare the morphology index datasets from the 2D and 3D analysis methods, soma area/volume and arborization area/convex hull volume were normalized on a scale from 0– 1 for each set of data. The morphology index was re-calculated using the normalized values and an unpaired T-test with Welch’s correction was used to directly compare the 2D and 3D datasets. These tests were performed in GraphPad Prism (v10.0.3 Graphstats Technologies).

To more accurately showcase the individual animal variability within each group, we also performed a nested 2-way ANOVA, with the animals nested within stress and stress within diet. This analysis was not performed by González Ibáñez et al. (2023). Due to this, we have re-analyzed the raw 2D microglia morphology data using a nested design. To accomplish this, we used a mixed-effects model, where diet and stress are fixed effects and animal is a random effect. We then computed the estimated marginal means for the combinations between diet and stress before performing pairwise comparisons between these means. The Tukey’s *post-hoc* test was applied to perform the pairwise comparisons, to allow for comparison with the non-nested data. The Kenward-Roger degrees-of-freedom method was used to estimate the degrees of freedom. This analysis was performed in RStudio v2024.04.2+764.

For all analyses, statistical significance was reached when *p* value ≤ 0.05 and potential changes were indicated at *p* < 0.1. All graphs were created in GraphPad Prism (v10.0.3 Graphstats Technologies).

## Results

### Direct 2D *vs* 3D Comparison

We first sought to perform morphological analyses that were most similar to those performed in González Ibánez et al. (2023) for direct comparison between the 2D and 3D morphology analysis methods. These analyses were performed for all cellular values within each group. For ease of reference, we will break up this section per morphological characteristic. Please refer to Table 2 for a comparison between different analysis methods and statistical designs in terms of statistical differences.

**Table 2.** |. Comparison of statistical outcomes for statistically significant differences across different analysis methods and statistical designs. n=3–5 mice/group; n=32–94 microglia/group. Statistical significance was assessed using a 2-way or nested 2-way ANOVA with Tukey *post hoc* test, where *p*≤0.05 is considered significant.

| Characteristic | Interaction | 3D | 3D % Change | Nested 3D | 2D | 2D % Change | Nested 2D |
| --- | --- | --- | --- | --- | --- | --- | --- |
| *Soma* | CD CTRL vs. KD CTRL | ns<br>0.12 | 89.90 | ns<br>0.81 | **0.02** | 86.81 | ns<br>0.17 |
|  | CD CTRL vs. CD SUS | **0.0004** | **84.41** | ns<br>0.44 | **0.0002** | **52.19** | **0.03** |
|  | CD RES vs. KD RES | **0.05** | **115.36** | ns<br>0.48 | ns<br>0.98 | 103.15 | ns<br>0.99 |
|  | KD CTRL vs. KD SUS | **0.02** | **87.70** | ns<br>0.73 | ns<br>0.99 | 98.09 | ns<br>0.99 |
|  | KD CTRL vs. KD RES | **0.01** | **113.94** | ns<br>0.49 | ns<br>0.51 | 106.77 | ns<br>0.75 |
|  | KD SUS vs. KD RES | **<0.0001** | **129.92** | ns<br>0.07 | ns<br>0.33 | 108.85 | ns<br>0.59 |
| *Branch Spread* | CD CTRL vs. KD CTRL | **<0.0001** | **61.00** | ns<br>0.16 | **0.04** | **85.12** | ns<br>0.66 |
|  | CD CTRL vs. CD SUS | **<0.0001** | **59.00** | ns<br>0.16 | ns<br>0.33 | 88.38 | ns<br>0.91 |
|  | CD CTRL vs. CD RES | ns<br>0.91 | 85.70 | ns<br>0.99 | **0.002** | **80.28** | ns<br>0.39 |
|  | CD SUS vs. CD RES | **0.007** | **145.25** | ns<br>0.42 | ns<br>0.50 | 90.83 | ns<br>0.92 |
|  | CD RES vs. KD RES | **0.01** | **69.54** | ns<br>0.52 | ns<br>0.73 | 106.85 | ns<br>0.99 |
| *Morphology Index* | CD SUS vs. KD SUS | **0.005** | **53.13** | ns<br>0.93 | ns<br>0.99 | 97.14 | ns<br>0.99 |
|  | KD CTRL vs. KD SUS | **0.0001** | **40.34** | ns<br>0.71 | ns<br>0.10 | 84.94 | ns<br>0.68 |
|  | KD SUS vs. KD RES | **<0.0001** | **283.87** | ns<br>0.23 | **0.007** | **124.60** | ns<br>0.36 |
| *Total Sholl Intersections* | CD CTRL vs. KD CTRL | **0.0006** | **75.98** | ns<br>0.59 | NA | NA | NA |
|  | KD CTRL vs. KD SUS | **<0.0001** | **151.57** | ns<br>0.40 | NA | NA | NA |
|  | KD SUS vs. KD RES | **0.005** | **74.28** | ns<br>0.78 | NA | NA | NA |
| *Critical Radius* | ns | ns | ns | ns | NA | NA | NA |

CD=Control diet, KD=Ketogenic diet, CTRL=control stress, SUS=susceptible phenotype to stress, RES=resistant phenotype to stress, ns=not significant, NA=not applicable. Numerical values are *p*-values and percent change values.

#### Soma Area vs Soma Volume

In terms of interactions, González Ibáñez et al. (2023) reported a significant interaction of stress x diet (*F*_(2,374)_=5.63, *p*=0.004) and a main effect of stress (*F*_(2,374)_=9.63, *p*=0.0009) for soma area. Our 3D analysis supports this, with a significant interaction observed for stress x diet (*F*_(2,339)_=8.32, *p*=0.0003) and a main effect of stress (*F*_(2,339)_=17.39, *p*=<0.0001).

Further investigation between groups revealed a significant difference (*p*=0.02) between the CD CTRL and KD CTRL groups, indicating larger soma area in the KD CTRL group in the 2D analysis. However, in our 3D analysis, we did not observe a significant difference between these groups (*p*=0.12) (Fig. 2B,E). In addition, González Ibáñez et al. (2023) reported a significant difference (*p*=0.0002) between the CD CTRL and CD SUS groups, with microglia from the CD SUS having a larger soma area. Our 3D analysis supports this finding, with an observed significant difference (*p*=0.0004) between these groups (Fig. 2B,E). In addition, our 3D analysis also found significant differences between KD CTRL and KD SUS (*p*=0.02) (larger soma for KD SUS), KD CTRL and KD RES (*p*=0.01) (larger soma for KD CTRL), KD SUS and KD RES (*p*=<0.0001) (larger soma for KD SUS), and CD RES and KD RES (*p*=0.05) (larger soma for CD RES) groups (Fig. 2B,E) that were not observed in the 2D analysis (*p*=0.99, *p*=0.51, *p*=0.33 and *p*=0.98, respectively). This highlights a potential difference between 2D and 3D analysis when it comes ascertaining differences in soma volumes.

#### Arborization Area vs Convex Hull Volume

For arborization area, González Ibánez et al. (2023) reported a significant interaction of stress x diet (*F*_(2,374)_=5.42, *p*=0.005) and a main effect of stress (*F*_(2,374)_=4.61, *p*=0.01). Similarly, our 3D analysis revealed a significant interaction between stress x diet (*F*_(2,339)_=15.00, *p*=<0.0001), but we did not observe a main effect of stress (*F*_(2,339)_=2.17, *p*=0.12). However, we did observe a significant main effect of diet (*F*_(1,339)_=10.61, *p*=0.001).

Similar to the soma area, the 2D analysis resulted in a significant difference between CD CTRL and KD CTRL (*p*=0.04) groups, with a larger arborization area in the KD CTRL group. This finding was supported by our 3D analysis (*p*=<0.0001) (Fig. 2D,F). The 2D analysis also resulted in a significant difference between CD CTRL and CD RES (*p*=0.002) groups, with CD RES having a higher arborization area, but this finding was not found by our 3D analysis (*p*=0.91) (Fig. 2D,F). In addition to these, our 3D analysis revealed significant differences between CD CTRL and CD SUS (*p*=<0.0001) (larger convex hull for CD SUS), CD SUS and CD RES (*p*=0.007) (larger convex hull for CD SUS), and CD RES and KD RES (*p*=0.01) (larger convex hull for KD RES) groups (Fig. 2D,F). These differences were not observed in the 2D analysis (*p*=0.33, *p*=0.50 and *p*=0.73, respectively). Similar to the soma data, this indicates a potential difference between the 2D and 3D analysis in terms of branch spread (i.e., arborization area and convex hull volume).

#### Morphology Index

González Ibáñez et al. (2023) reported a significant main effect of stress (*F*_(2,374)_=9.63, *p*=<0.0001) for the morphology index. Our 3D analysis supports this finding (*F*_(2,339)_=6.94, *p*=0.001), and further revealed a significant interaction of stress x diet (*F*_(2,339)_=8.27, *p*=0.0003).

Further analysis revealed a significant difference between KD SUS and KD RES groups in the 2D (*p*=0.007) and 3D (*p*=<0.0001) analysis, with the KD SUS group having a higher value (Fig. 2G). Our 3D analysis further revealed significant differences between KD CTRL and KD SUS (*p*=0.0001) (higher value for KD SUS), as well as CD SUS and KD SUS (*p*=0.005) (higher value for KD SUS) groups (Fig. 2G). These results were not observed in the 2D analysis (*p*=0.10 and *p*=0.99, respectively).

Overall, these results suggest that although similar, the 2D and 3D analysis methods do result in distinct differences between the analyzed groups, with often more differences observed in the 3D analysis method. These differences observed with the 3D method are also accompanied by a greater percent change (Table 2).

### Further 3D Morphological Characterization

In addition to soma volume, convex hull volume and morphology index, we quantified the number of Sholl intersections at 5 μm radius intervals, and the critical radius (i.e., the radius with the highest number of intersections). Although these measurements are possible to quantify in 2D, they were not performed in González Ibáñez et al. (2023).

The critical radius analysis did not indicate any significant differences between groups (Fig. 2I). However, the Sholl analysis revealed a significant interaction between stress x diet (*F*_(2,339)_=6.39, *p*=0.002), and main effects of stress (*F*_(2,339)_=10.29, *p*=<0.0001) and diet (*F*_(2,339)_=6.61, *p*=0.01). Our *post-hoc* analysis indicated that there is a significant difference between CD CTRL and KD CTRL (*p*=0.0006), KD CTRL and KD SUS (*p*=<0.0001), as well as KD SUS and KD RES (*p*=0.005) groups (Fig. 2C,H). Among these differences, KD CTRL and KD RES both have a greater number of total Sholl intersections compared to their respective comparisons, perhaps indicating an increased number of branches or more complex branching patterns in these groups.

### Exploration of a Nested Statistical Design

We next sought to investigate how the 2D and 3D microglia morphology analysis methods could differ with the utilization of a different statistical method that also takes into account individual animal variability. As a note, the data reported in González Ibáñez et al. (2023) used individual cell values as a sample size to examine the variability of microglial morphology between cells, and did not use animal means which do not take into account intercellular variability. However, the nested design takes into account both cellular and animal variability. Therefore, we performed a nested 2-way ANOVA in RStudio with the raw 2D microglia morphology data from González Ibáñez et al. (2023) and re-analyzed data from the 3D microglia morphology method.

#### Soma Area vs. Soma Volume

The 2D data analyzed with a nested 2-way ANOVA revealed a significant interaction of stress x diet (*F*_(2,374)_=5.76, *p*=0.004) and a main effect of stress (*F*_(2,374)_=6.97, *p*=0.001), which is supported by the 3D data (stress x diet (*F*_(2,339)_=9.66, *p*=<0.0001); stress (*F*_(2,339)_=22.28, *p*=<0.0001)).

Tukey’s *post-hoc* indicated that there was a significant difference between CD CTRL and CD SUS (*p*=0.003) groups in the 2D analysis (Fig. 3Aii), with CD SUS having larger soma area. We did not observe any significant differences between groups in the 3D analysis (Fig. 3Ai). However, we observed a potential increase in soma volume in the KD SUS group, compared to the KD RES (*p*=0.07) group in the 3D analysis (Fig. 3Ai).

**Figure 3.**
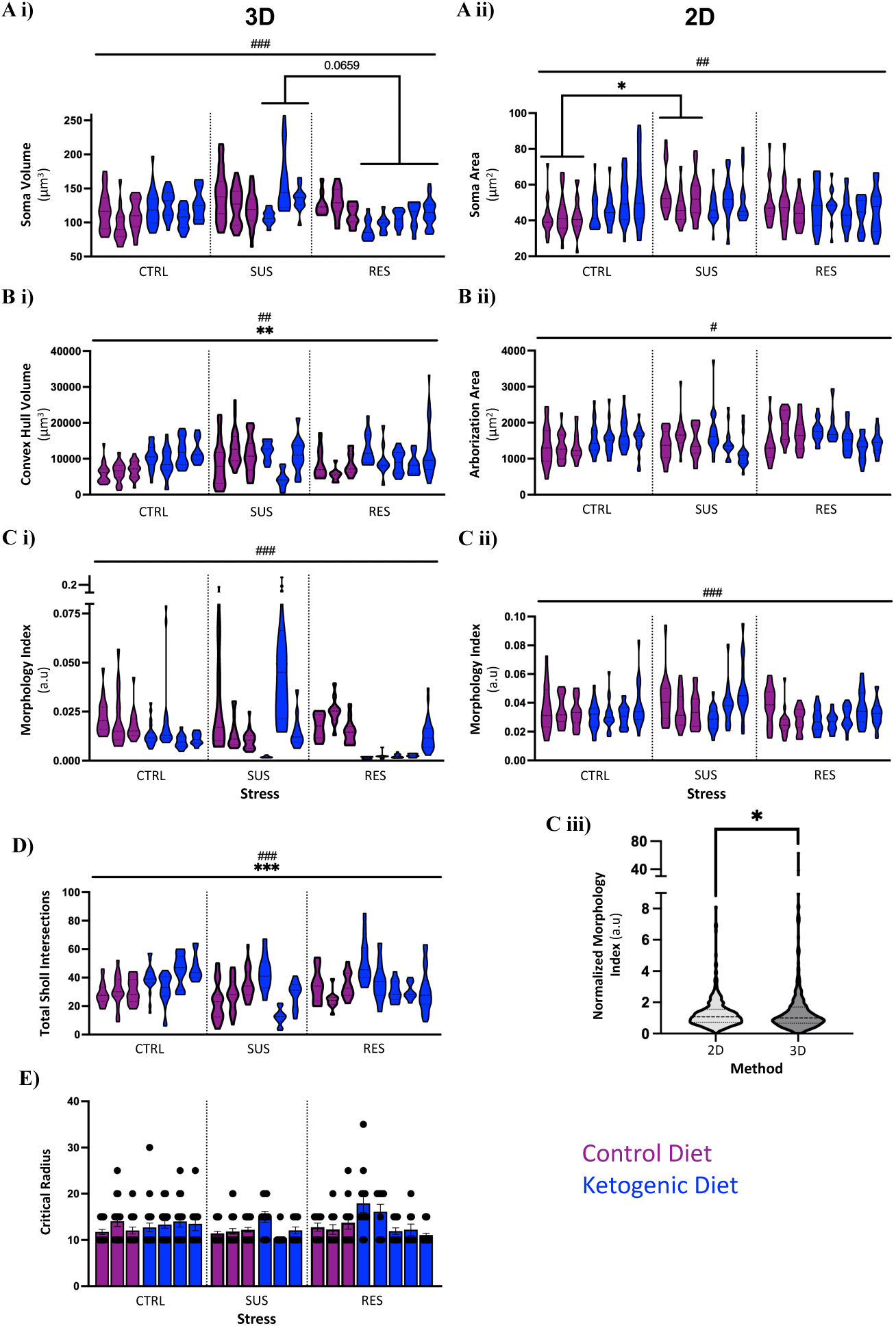
| Comparison between the 2D and 3D microglial morphology analysis methods using a nested 2-way ANOVA. Significant differences were found for soma area where the CD SUS group has a larger soma area compared to CD CTRL (Aii). No other comparison resulted in significant differences in either the 2D or 3D morphology analysis method with this statistical method. However, when the morphology index was calculated with normalized values, a significant difference was observed when the entire datasets were compared to one another (Ciii). n=3–5 mice/group; n=32–94 microglia/group. Statistical significance in Ciii was assessed using an unpaired t-test with Welch’s correction. All other tests consisted of a nested 2-way ANOVA with Tukey’s *post hoc* test and data are expressed as mean ± standard error of the mean, where \**p*≤0.05, \*\**p*<0.01, \*\*\**p*<0.001, *\*\*\*\*p*<0.0001. #≤0.05, ##<0.01, ###<0.001 depicts a main effect of stress. CD: control diet; KD: ketogenic diet; CTRL: non-stressed controls; SUS: susceptible to stress; RES: resistant to stress; a.u: arbitrary units.

#### Arborization Area vs. Convex Hull Volume

For the 2D arborization area, a significant interaction was observed for stress x diet (*F*_(2,374)_=6.09, *p*=0.003), as well as a main effect for stress (*F*_(2,374)_=3.76, *p*=0.02). The 3D analysis for convex hull volume resulted in a significant interaction of stress x diet (*F*_(2,339)_=16.94, *p*=<0.0001), as well as main effects of stress (*F*_(339)_=5.03, *p*=0.007) and diet (*F*_(2,339)_=9.21, *p*=0.003). However, we observed no significant differences after the *post-hoc* test for either analysis method (Fig. 3Bi,ii).

#### Morphology Index

The nested analysis revealed significant main effects of stress for both the 2D (*F*_(2,374)_=10.75, *p*=<0.0001) and 3D (*F*_(2,339)_=9.06, *p*=0.0001) analysis methods. Additionally, a significant interaction was observed in the 3D analysis between stress x diet (*F*_(2,339)_=11.13, *p*=<0.0001). We observed no significant differences after the *post-hoc* test for either analysis method (Fig. 3Ci,ii). We also qualitatively observed a tendency for morphology index values to be higher when generated using the 2D analysis method compared to the 3D analysis method. To investigate this further, we normalized the data from both analysis methods and directly compared the entire datasets using normalized morphology index values through an unpaired T-test with Welch’s correction. We observed a significant difference (*p*=0.05) between the 2D and 3D methods, with the 3D method generating lower overall values compared to the 2D method.

#### 3D Sholl Analysis

We observed a significant interaction between stress and diet (*F*_(2,339)_=8.69, *p*=0.0002), as well as main effects of stress (*F*_(2,339)_=11.28, *p*=<0.0001) and diet (*F*_(2,339)_=20.91, *p*=<0.0001) for the 3D Sholl analysis. However, there were no significant differences between groups after our *post-hoc* analysis (Fig. 3D). No statistically significant effects were found for critical radius in the 3D dataset (Fig. 3E).

Overall, when statistical tests more accurately account for individual animal variability within the groups, many of the statistically significant comparisons between individual groups using whole group data are not observed, especially for the 3D dataset.

## Discussion

This methods research paper set out to compare 2D and 3D morphology analysis tools of IBA1^+^/TMEM119^+^ microglia from the ventral hippocampus of adult male mice imaged using a confocal microscope (40x) from a previously published research paper from our group, González Ibáñez et al. (2023). The 2D microglia morphology method used in González Ibáñez et al. (2023) is a well-established methodology that has been utilized in multiple publications over the past decade (Basilico et al., 2022; Bordeleau et al., 2020, 2021; Chen et al., 2024; Enlow et al., 2021; Fernández De Cossío et al., 2021; González Ibáñez et al., 2023; Hui et al., 2019; Hui, St-Pierre, Detuncq, et al., 2018; Hui, St-Pierre, El Hajj, et al., 2018; Lecours et al., 2020; Paasila et al., 2019; Picard et al., 2021; Savage et al., 2020; Tremblay et al., 2012). The 3D microglial morphology method used in this study was performed using Imaris v10.0.2, a well-known and powerful analysis software for 3D microscopy data (Haass-Koffler et al., 2012; Nguyen & Thompson-Peer, 2021; Rose et al., 2024; Singaravelu et al., 2017; Testen et al., 2024; VanRyzin et al., 2019). Although microglial/glial cell morphology analysis has not been extensively performed using Imaris in comparison to 2D methods, its potential—based on the available options within the Imaris workspace—is notable.

Overall, we found measurable differences between the two methods, with the 3D analysis method producing overlapping, but also distinct significant differences between groups to a greater extent than with the 2D analysis. In addition, a nested statistical design—which more accurately takes into account individual animal variability—produced distinct results compared to total cell/group data, which perhaps highlights more robust findings.

### 2D vs 3D Microglia Morphology Analysis

The most striking differences between the 2D and 3D methods were related to the soma. Our previous 2D analysis by González Ibáñez et al. (2023) indicated two statistically different specific comparisons between groups, whereas the 3D analysis indicated five. However, the differences observed between CD CTRL and KD CTRL in the 2D analysis were not found in the 3D analysis. It is not surprising that soma should produce the most varied responses between these methods. The 2D analysis involves a Z-stack image that is converted to a maximum intensity projection image, resulting in a soma area that is equal to the widest portion of the soma, whereas the 3D analysis takes into account soma directionality in all three planes, as well as indentations and inconsistencies in soma shape. These can be common in microglia, and many other cells, especially when in apposition to other cells or structures, such as blood vessels or axons (Stratoulias et al., 2019). Therefore, it is possible that the 3D analysis is better able to capture slight variations in soma shape, revealing changes to cell populations within groups that the 2D analysis method cannot.

A similar argument can also be made for arborization area. Our 3D analysis indicated significant differences in arborization area (measured by convex hull volume) between CD CTRL and KD CTRL groups, a finding which has been observed by our group and others (González Ibáñez et al., 2023; Gzielo et al., 2019). However, there were also distinct interactions found in 2D that the 3D analysis did not capture, and *vice versa*. The most likely cause of this is the fact that although microglia maintain distinct territories, microglial branches do somewhat overlap in 3D space, and microglial clusters can form during an insult (e.g., lesions in multiple sclerosis), as well as in health (Hume et al., 1983; Van Horssen et al., 2012). The maximum projection images make it extremely challenging to differentiate between overlapping branches, resulting in slightly less accurate depictions of arborization area. This being said, the 2D analysis method is by no means considered incorrect, but is instead another method with strengths and limitations that should be contemplated before performing analysis. This is also the case with 3D analysis. Although differentiating between overlapping branches is relatively simple, 3D analysis software (*e.g.*, Imaris) does not always accurately depict branch length. For example, a recent study comparing different morphology analysis tools for neurons of the *Drosophila* larvae reported that although automatic detection using 3D analysis software (*e.g.*, Imaris) was the most efficient and convenient method for dendrite reconstruction of the various methods tested, automatic detection of dendrites in Imaris resulted in significantly shorter dendritic lengths compared to manually traced dendrites (Nguyen & Thompson-Peer, 2021). However, they also reported that this is somewhat mitigated by the ability to easily edit reconstructed objects manually, a feature that we also relied upon in our analysis. It is also important to note that the 2D microglial morphology analysis has a similar number of possible quantifiable measurements compared to the 3D analysis, further bolstering its capabilities (Leyh et al., 2021).

Although the 3D method may be able to pick up on more subtle changes to morphology, such as soma indentations, and, by extension, between groups, robust morphological changes are still captured by the 2D analysis. For example, the morphology index—a more descriptive measure of changing microglial morphology as it takes into account both soma size and arborization area— was observed to be significantly higher in the KD SUS group compared to the KD RES group in both the 2D and 3D analysis methods (Fig. 2G).

Nonetheless, despite the noticeable differences between these two methods, the overall findings of González Ibáñez et al. (2023) for microglial morphology using 2D analysis methods are supported by the 3D analysis method. Ketogenic diet and stress do have an effect on microglial morphology at baseline, and based on the 3D morphological data, the findings reported by González Ibáñez et al. (2023)—that ketogenic diet can help prevent some of the morphological adaptations microglia have in response to repeated social defeat stress—are supported. Similar findings between 2D and 3D methods with respect to microglial morphology have also been found by others. For example, microglia process length was found to be significantly shorter in the inferior temporal cortex from post-mortem human brain tissue with Alzheimer’s disease pathology compared to age-matched controls using ImageJ (2D) and Imaris software (3D) (Davies et al., 2017).

#### Important Considerations for Method Choice

Although the most important consideration for choosing an analysis method is arguably the research question at hand, varied financial and temporal constraints always accompany each method and influence choice. The 2D analysis performed by González Ibáñez et al. (2023) was performed using ImageJ, a free open-source software funded by the National Institutes of Health (NIH). This software has a wide-array of easily installed plugins that can perform an impressive number of analyses for cellular morphology and much more. Another free open-source software that can be used for cellular morphology is QuPath. QuPath is extremely user-friendly and has some built-in analytical features, such as cell detection. These programs can also be combined, where more easily made annotations performed in QuPath can be sent to ImageJ through QuPath, and subsequently analyzed using the very convenient ImageJ plugins. Additionally, users can enter commands and write scripts in both QuPath and ImageJ, allowing for customizable workflows that can drastically enhance analysis speed and contribute to reproducibility.

Although Z-stacks can be imported into both QuPath and ImageJ—as well as the fact that ImageJ has some 3D analysis plugins—these programs are quite limited in 3D at this time. This being said, recent years have seen a surge in interest in 3D morphology characterization, and open-source plugins for ImageJ (e.g., GIANI), as well as other programs such as Microscopy Image Browser, are becoming more powerful and convenient for use in 3D (Barry et al., 2022; Belevich et al., 2016). However, in terms of time, these options still take decidedly longer to analyze cells in 3D compared to software built for 3D analysis, like Imaris, especially considering batch analysis capabilities, where analysis parameters can be applied to multiple images at a time. For example, manually tracing neurons from the *Drosophila* larvae took an average of 21 minutes per image, whereas Imaris had an average tracing time of 7 minutes per image (Nguyen & Thompson-Peer, 2021).

Imaris is an incredibly powerful tool for visualization and analysis of 3D microscopy data. Built-in plugins are available in the software, but custom plugins can also be mapped and performed in Imaris. For example, this project utilized a plugin built by Ironhorse1618 (2023): https://github.com/Ironhorse1618/Python3.7-Imaris-XTensions, which extracts and sends reconstructed objects from Imaris to Python 3.7. Subsequent calculations (i.e., soma volume, Sholl analysis, convex hull, etc.) are performed in Python before the data is sent back to Imaris for ease of visualization and use. Overall, software like Imaris is extremely powerful, versatile and efficient, but also financially restrictive. The upfront costs for software such as these are significant, and are often accompanied by yearly subscription fees. However, more affordable options through core imaging facilities, that in some cases can be accessed remotely, may exist.

Overall, the decision to use either 2D or 3D analysis methods for cell morphology depends heavily on the research question, but also on financial constraints, time and experience of the researcher, as was also highlighted by Green et al. (2022). In Green et al. (2022), inconsistencies between ImageJ-based methods for quantifying microglial morphology in 2D were highlighted. It was suggested that combining methods would provide the most accurate depiction of microglial morphology, as was previously discussed in this paper (Green et al., 2022). The authors also emphasized that single-cell analyses are preferable to aggregate data calculated from the means of many microglia within the same photomicrograph (Green et al., 2022). We agree with this suggestion, as means compiled from multiple biological replicates may mask natural variability within the microglial population in terms of morphology. However, single-cell analyses may also result in certain biological replicates driving the dataset—since animal variability is common— and somewhat arbitrarily increase the sample size (Voelkl et al., 2020). To account for this, a nested or Bayesian statistical design can be utilized (Coventry & Bartlett, 2024; Galbraith et al., 2010).

Our study revealed differences between data analyzed with a nested 2-way ANOVA with ‘animal’ as our random factor *vs* a 2-way ANOVA. Both the 2D and 3D datasets analyzed with the nested statistical design resulted in much fewer significant differences between groups. The 3D data in particular was drastically different, as no statistically significant comparisons between individual groups were maintained using the nested design. A likely reason for this is that the pooled individual data points for each group used in the 2-way ANOVA assumes they are all independent from one another, and does not account for similarities within each individual animal, ultimately resulting in an increase in Type 1 errors (Galbraith et al., 2010). Therefore, the nested approach potentially yields more accurate and robust statistically significant effects when compared to pooled group data used in the 2-way ANOVA. However, it is important to note that the Tukey *post-hoc* test is a particularly stringent test, as it makes all possible comparisons despite their biological/experimental relevance, and adjusts *p*-values to account for this. A *post-hoc* test that computes multiple comparisons based off of selected comparisons (e.g., Šídák or Bonferonni multiple comparisons tests) that are biologically and experimentally relevant may be more appropriate. Nonetheless, our goal was to compare the non-nested 2-way ANOVA with a nested 2-way ANOVA, meaning that the *post-hoc* test, as long as it is consistent between the two designs, is somewhat irrelevant.

Overall, we believe that this methods paper will help shed new light on the differences between 2D and 3D microglial morphology analysis. We also aim to stress that although outcomes are different, general tendencies and overall findings are consistent between the two, and therefore no one method is considered correct or incorrect. Instead, the right method of choice should depend on the research question and whether or not it can be sufficiently and accurately answered using 2D or 3D image analysis tools.

## Conclusion

We re-analyzed raw confocal microscopy images of microglia in 3D from a previously published paper from our group that analyzed microglia morphology in 2D in the ventral hippocampus of adult mice. Overall, the 3D analysis method supports many of the observations made using the 2D analysis method, but also indicates many significant differences between groups that were not observed using the 2D method. This suggests that the 3D analysis method may be better able to capture minute changes in microglial morphology, possibly due to superior visualization of the cell compared to collapsed 2D images. However, overall conclusions about microglial morphological adaptation to ketogenic diet and stress from both methods are similar, highlighting the fact that both methods are appropriate for this type of analysis. In addition, we compare data analyzed using a nested and non-nested design, showing how individual animal variability can influence datasets.

## Acknowledgments

We acknowledge with respect the ləkʷəŋən peoples on whose traditional territory the University of Victoria stands and the Songhees, Esquimalt and WSÁNEĆ peoples whose historical relationships with the land continue to this day. We also acknowledge the animal work done by Drs. Kaushik Sharma, Nathalie Vernoux and Kanchan Bisht, who are authors on the original manuscript. In addition, we acknowledge that image analysis for this work was performed in the University of British Columbia Life Sciences Institute Imaging Core Facility, RRID: SCR_023783. We also thank the Imaris application support team for their guidance for the use of Imaris software.

CJM was supported by a CIHR CGS-M scholarship, and a graduate grant from the Branch Out Neurological Foundation. EDT was funded by the KF Hein grant, Jo Kolk grant, Hendrik Muller grant, and the UMC Utrecht Strategic Network Development grant. HAV was a CIHR Fellow, Michael Smith Health Research BC Research Trainee and was supported by a Brain Canada Training Fellowship and BC Women’s Health Research Institute Fellowship. FGI is a Michael Smith Health Research BC research trainee and was supported by a full doctoral scholarship by the Mexican Council of Science and Technology (CONACYT, now CONAHCYT). M-ÈT is a Tier II Canada Research Chair in *Neurobiology of Aging and Cognition* and holds funding through the Natural Sciences and Engineering Research Council of Canada (RGPIN: 2024-06043), Canadian Institutes of Health Research (CIHR) (PJT461831) and the Canadian Foundation for Innovation (John R. Evans Leaders Fund: 39965).

## Data Availability Statement

The data that support the findings of this study are available from the corresponding author (M-ÈT) upon reasonable request.

